# A Zika virus-associated microcephaly case with background exposure to STORCH agents

**DOI:** 10.1101/052340

**Authors:** Mauro Mitsuru Hanaoka, Alexandre Fligelman Kanas, Carla Torres Braconi, Érica Araújo Mendes, Robert Andreata Santos, Luís Carlos de Souza Ferreira, Marielton dos Passos Cunha, Patrícia Beltrão Braga, João Leonardo Mendonça Dias, Carolina Manganeli Polonio, David Anibal Garrido Andrade, Carla Longo de Freitas, Cristiano Rossato, Wesley Nogueira Brandão, Jean Pierre Schatzmann Peron, Antonio Gomes de Amorim Filho, Joelma Queiroz Andrade, Rossana Pulcineli Vieira Francisco, Fernando Kok, Lisa Suzuki, Claudia da Costa Leite, Leandro Tavares Lucato, Amadou Alpha Sall, Paolo Marinho de Andrade Zanotto

## Abstract

We present a case of microcephaly associated with Zika virus (ZIKV) in a chronological, multimodal imaging approach, illustrating the hallmarks of this disease on intrauterine morphological ultrasound, transfontanelar ultrasound, computed tomography (CT) and magnetic resonance imaging (MRI). We also determined the serological e immunological status of the mother and newborn. Noticeably, there was evidence for maternal infection by ZIKV, cytomegalovirus (CMV), herpes simplex virus (HSV), dengue virus (DENV) and *Toxoplasma gondii*, which indicates a possible role of previous exposures to STORCH agents and possibly comorbidities in the severe fetal congenital manifestation.

**Author Summary:** Zika virus (ZIKV) is an emerging mosquito-borne arbovirus causing dengue-like symptoms. In humans the illness is characterized by malaise and cutaneous rash and absent or short-termed fever. Recently, the Brazilian Ministry of Health reported an outbreak of microcephaly in Brazil as a delayed effect of the 2014-2015 outbreak of ZIKV in the Northeast of Brazil. A 20-fold increase in the notifications of newborns with microcephaly was documented during the second semester of 2015. This increase was almost immediately found to be associated with ZIKV infections, both in Brazil and, retrospectively, in French Polynesia. Herein we report a case of microcephaly associated with ZIKV and we also present evidence of other maternal infections. Our results indicated that, both mother and microcephaly infant had immunologic status compatible with previous exposure (in the mother) by STORCH agents. These indicate a possible role of previous exposures and possibly comorbidities in the severe fetal congenital manifestation. □

## Introduction

Zika virus (ZIKV), a flavivirus, was isolated in 1947 in Uganda [1]. The virus is vectored by several *Aedes* mosquitoes [2]. ZIKV, caused sporadic human infections until 2007, when it caused an epidemics in Micronesia [3]. Latter in 2013, it caused an outbreak in Tahiti, French Polynesia and later in New Caledonia and the Cook Islands in 2014 [4]. It arrived in the Americas possibly in 2013-14 [5,6]. ZIKV is now causing outbreaks in Brazil, and several other countries in south and Central America. Cases started to be reported in in the Northeast of Brazil during March 2015, and the virus has been reported in most States in Brazil [7]. Importantly, a 20-fold increase in the notifications of newborns with microcephaly was documented during the second semester of 2015 [8]. In 2015, the Health authorities in Brazil reported an increase in the incidence of cases of microcephaly that was then assumed to be a delayed effect of the 2014-2015 outbreak of Zika virus (ZIKV) in the Northeast of Brazil. Microcephaly was then found to be associated with ZIKV infections, both in Brazil and, retrospectively, in French Polynesia. Later as the epidemic unfolded, the World Health Organization (WHO) declared ZIKV as a global health emergency in February 2016. The association of ZIKV and microcephaly was further confirmed by positive detection of the virus by RT-PCR only in the fetal brain sample [9]. More recently, arthrogryposis and several degrees of neuropathology, including encephalopathies, have been observed to associate with ZIKV infection. Herein we report a microcephaly case diagnosed by prenatal and postnatal imaging with evidence for maternal infection by ZIKV with a background of exposure to other pathogens.

## Materials and Methods

### Ethics Statement

We report a case from our quaternary care hospital. All patients received routine standard medical care. They were not submitted in any way to neither experimental exams nor treatment. This study was approved by our Institutional Review Board (IRB) and written informed consent, from the patient, was obtained for the publication of the images in the article. The research was approved by the Ethics Committee on the Research with Humans (CEPSH - Off.011616) of the Institute of Biosciences of the University of São Paulo.

### Ultrasound

Obstetric images were acquired with a GE Voluson E8 and transfontanelar images were obtained with a GE Logiq E9 (GE Healthcare, Pittsburg, PA, USA).

### Computed tomography (CT)

Images were acquired with a Brilliance 64 CT (GE Healthcare, Pittsburg, PA, USA) using a reduced radiation exposure protocol, compliant with the Alliance of Radiation Safety in Pediatric Imaging (The Image Gently Alliance) CT protocol recommendations. Post-processing of the CT data was performed with Philips iSite Radiology (Koninklijke Philips Electronics N.V., Netherlands).

### Magnetic Resonance Imaging (MRI)

Images were acquired Optima MR360 Advance (GE Healthcare, Pittsburg, PA, USA) without sedation of the patient.

### Real-time PCR

Samples of saliva, urine and total blood of the mother were taken to a level 2 laboratory for processing two months and 9 days after delivery. RNA was extracted from each sample using QIAamp UltraSens Virus Kit (Qiagen) or Trizol^®^ reagent (Invitrogen). Primers/probes used for ZIKV, Yellow fever (YF), Dengue virus (DENV) and Chikungunya (CHIKV) were previously described [3,10,11]. RPLA27 was used as endogenous control for the PCR reactions [12]. All assays were performed using the AgPath-IDTM One-Step RT-PCR reagents (Applied Biosystems).

### Virus strains

Control strains of dengue virus 2 NGC (DENV), Chikungunya S27 (CHIKV), Yellow fever virus (YFV-17D vaccine strain), Zika virus MR 766 (ZIKV) and West Nile virus (WNV) B956 lineage 2, from Africa were provided by the Institute Pasteur of Dakar, Senegal.

### Enzyme-linked immunosorbent assay (ELISA)

Viruses infections were determined by qualitative assays carried out with capture IgM and indirect IgG ELISA using a specific viral antigen for YFV, DENV, CHIKV and ZIKV, as previously described [13]. The titration of specific serum antibodies was made as previously described [14], with an antigen adaptation. Sera were individually quantified for the presence of specific antibodies directed for ZIKV and DENV2 NGC virus particles.

### Plaque reduction neutralization test (PRNT)

Because of the high degree of serologic cross-reactivity among flaviviruses, the seropositive samples were tested for the presence of neutralizing antibodies using a PRNT protocol. PRNT tests were made separately for West Nile virus, ZIKV and YFV, as previously described [15–17]. To evaluate the presence of neutralizing antibodies for DENV in the sera from the patients, we performed an *in vitro* neutralization test against DENV2 NGC strain as previously described [18]. Plaques were counted and percentages of antibody-dependent plaque reduction with regard to non-immune control sera were calculated. Neutralizing titers were expressed as the serum dilution yielding a 90% reduction in plaque number (PRNT_90_). Sera with PRNT_90_ were considered to have neutralizing antibody against West Nile, DENV, ZIKV and YFV. ELISA and PRNT experiments were done in duplicate and triplicate, respectively.

### Indirect Immunofluorescence assay

C6/36 cell line was previously infected with ZIKV. Serum from the mother and a negative control was diluted 1:10 and incubated at 37^o^C for 30 min, washed three times using PBS and incubated with anti-human IgG FITC-conjugated (Chemicon, Temecula, CA) at 37°C for 30 min. Slides were covered using glycerin and observed under fluorescent microscope (Olympus, Shinjuku, Tokyo, Japan).

### Flow cytometry

PBMCs were obtained by conventional Ficoll Hystopaque gradient centrifugation. 5x10^5^ cells were suspended in RPMI 10% FCS, plated in 24-well plates and stimulated or not with anti-CD3/CD28 (2 μg/ml) or phorbol-myristate acetate (PMA 50 ng/mL) + ionomycin (1μg/ml) plus Brefeldin A (1 μg/ml) for 12 hours. Further, samples were collected and submitted to flow cytometry staining protocols. Briefly, samples were incubated with fluorochrome-conjugated antibodies anti-CD4 APC and anti-CD8 PercP in 25 μl at a dilution of 1:50 at 4^o^C. Cells were washed twice in PBS and centrifuged at 450 g at 4^o^C for 5 min. For intracellular staining, samples were suspended in cytofix for 30 min and then added with 50 μl of permwash and centrifuged at 450 g at 4^o^C for 5 min. Samples were suspended in 25 μl of permwash containing fluorochrome conjugated antibodies anti-IFN-γ PE and anti-IL-17 FITC (BD Biosciences) and kept at 4°C. After incubation, all samples were washed twice in permwash and suspended in paraformaldehyde 1%. Data were acquired and analyzed using the Accuri C6 flow cytometer (BD Biosciences).

### Cytokine secretion

Supernatants collected from the samples plated, as for flow cytometry without brefeldin, were used for cytokine secretion evaluation by *cytometric bead array* (BD Biosciences), according to the manufacturer’s protocol.

## Results

### Brief case description

A 32-year-old woman and her child, from Santos, São Paulo State, Southeast of Brazil reported having had dengue fever in the first trimester of 2013, coinciding with a massive DENV-4 outbreak in Santos. Her last menstruation was on February 6th, 2015 and she had no recent travel history. Around 3-4 weeks of gestation (March, 2015), she developed a febrile illness that lasted a few days, presenting with low-grade fever; maculopapular exanthema; non-purulent conjunctivitis; transient bilateral and symmetric polyarthralgia, myalgia and asthenia. On April 5th, she started prenatal care in Santos and obstetric ultrasound was normal and confirmed the gestational age. She had a VDRL with a titer of 1:2 and a FTA-ABs IgG (+) and IgM (-). As consequence, she and her partner were treated for syphilis (caused by *Treponema pallidum*). After three doses of penicillin, her VDRL titer remained 1:2. She also had IgG reactive to CMV, HSV 1 and 2 and slightly above cutoff value to *T. gondii* and CMV (Table 1). Her rubella seropositivity was deemed irrelevant, since she was vaccinated on a previous gestation. On July 13th, at 22 weeks of gestation, she was diagnosed with an upper respiratory tract infection and received symptomatic treatment. On July 31st, morphological ultrasound showed minimal fetal pericardial effusion and microcephaly. At 32 weeks (09/21) of gestation, these findings were confirmed by the fetal medicine unit at USP and she was admitted to the quaternary care hospital. Additional obstetric ultrasounds were performed (Figure 1) and showed developmental delay and microcephaly. Moreover, hypoplastic nasal bone, retrognathia, absent cavum septum pellucidum, periventricular calcification and ventriculomegaly were also observed. Fetal echocardiography was unremarkable. Noticeably, serology tests (Table 1) confirmed that she had IgG for CMV and HSV and a 32 fold increase in the anti *T. gondii* IgG titer and a 2 fold increase in the anti rubella IgG titer and the mother was found to be HIV-1 negative. A karyotype test was also performed, which revealed a 46 XX pattern, ruling out chromosomal disorders. At 38 weeks of gestation (11/03), she had a normal Cesarean delivery, without complications with an APGAR score 9/10/10, weighting 2275g, 40.5 cm in length and with a head circumference of 26.5 cm (three standard deviations (SD) below the reference value for the child’s age). Congenital syphilis was ruled out because the child had a serum VDRL titer of 1:1 (inferior to the maternal titer of 1:2); negative cerebrospinal fluid (CSF) and urine for VDRL, unremarkable complete blood count (CBC) and whole body radiography (supplementary figure 1). Immunohistochemical analysis of the placenta revealed no antigens for syphilis, CMV, adenovirus and HSV at the time of birth. Serological testing, after 8 days after birth, indicated that the baby was IgG positive for *T. gondii*, HSV and CMV (Table 1). Because anti *T. gondii* IgG comes from the mother, the continuous titer increase in time could suggest that she may have experienced a reinfection or reactivation of toxoplasmosis during pregnancy.

**Table 1.**
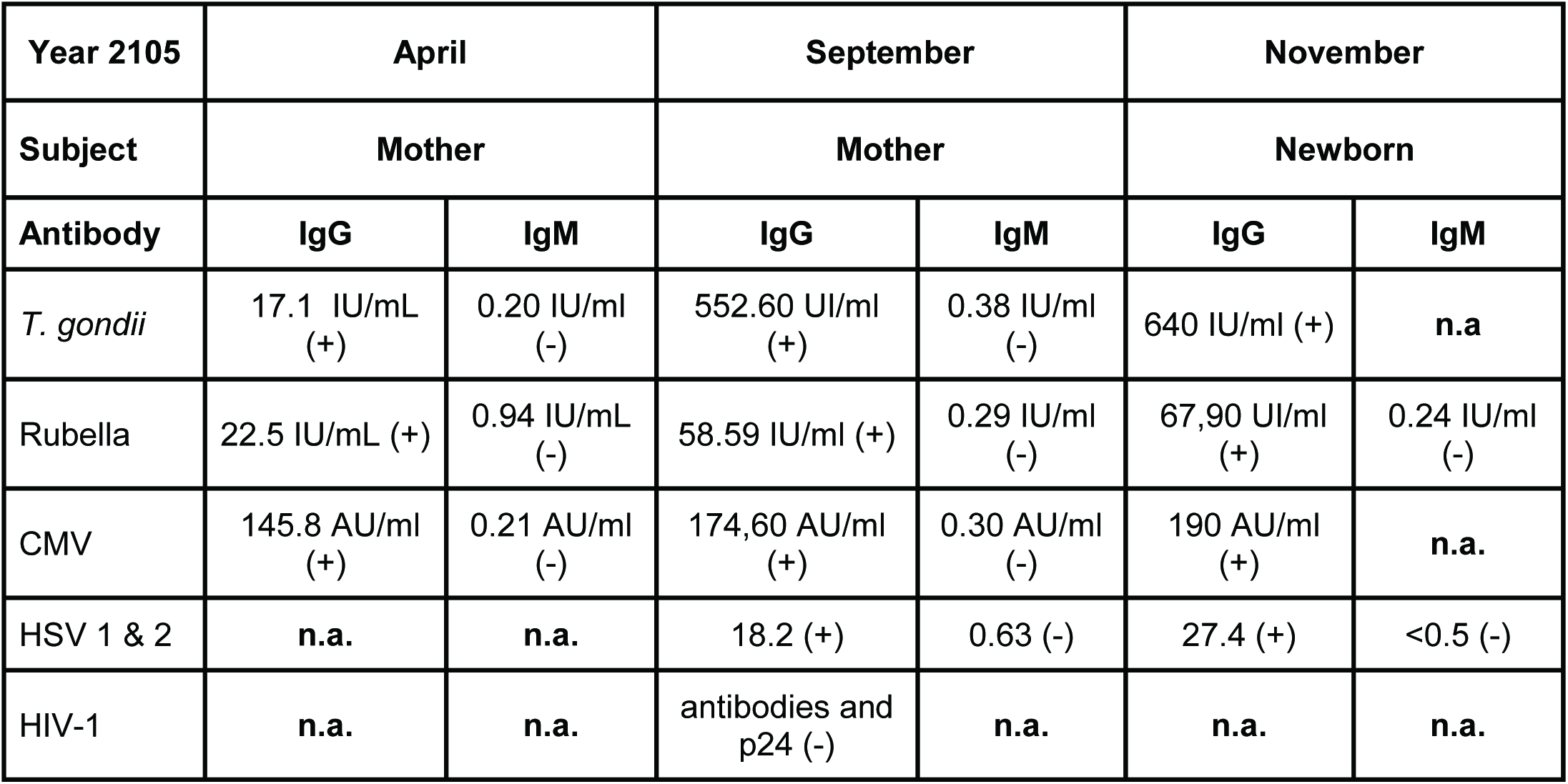
Serological clinical analysis. Routine laboratory exams in April were done at an outside facility (AFIP), in Santos. All other exams were done at USP Hospital Central laboratory. Although these exams follow strict validation rules imposed by health authorities, direct comparison of values from the two diagnostics units has to be made with care, since aspects of the protocols may differ.

**Figure 1.**
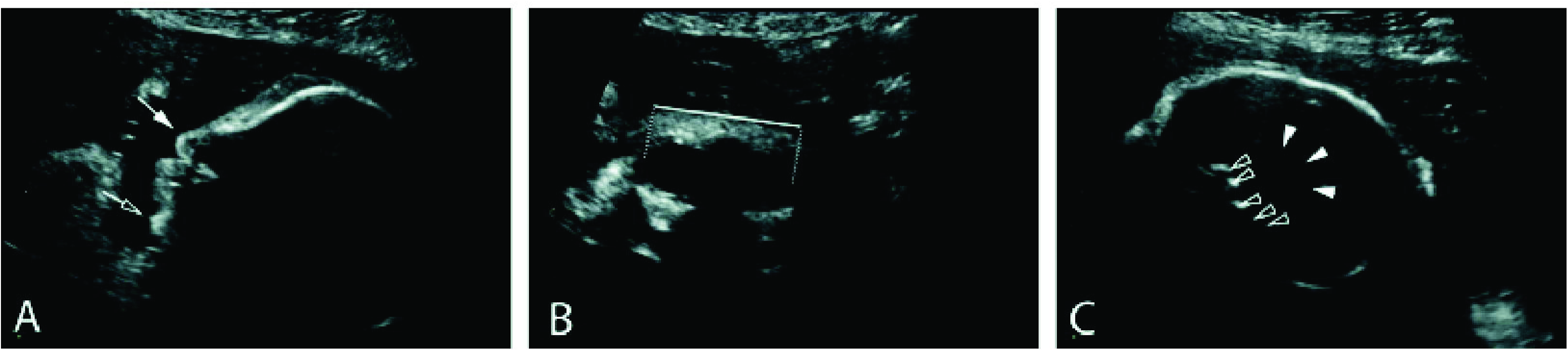
Second trimester obstetric ultrasound. Sagittal view of the fetal face (A) presents hypoplastic nasal bone (black arrow) and retrognathia (white arrow). Coronal view of the fetal face (B) discloses hypotelorism (white line). Parasagittal view of the fetal head (C), showing ventriculomegaly (white arrowheads) and periventricular calcifications (black arrowheads).

### Imaging findings

CT scans of the newborn’s head (Figure 2) presented a pronounced microcephaly, volumetric reduction of brain parenchyma with a simplified gyral pattern and shallow Sylvian fissures. There were also cortical, subcortical, basal ganglia and coarse periventricular calcifications with increased supratentorial cerebrospinal fluid spaces. No abnormalities were identified in the posterior fossa. 3D reconstruction images (Figure 2 D and E) show occipital protrusion and overriding parietooccipital bones. Transfontanellar ultrasonography at 3 months of age also showed volumetric reduction of the brain, simplified gyral pattern, asymmetric ventriculomegaly and cortical, subcortical and basal ganglia calcifications (Figure 3) albeit some further development of brain could be noticed. Magnetic resonance image (MRI) showed a craniocephalic disproportion with volumetric reduction of brain parenchyma, characterizing severe microcephaly and deformity of the skull, with occipital protrusion and overriding of parieto-occipital bones, cortical dystrophy, poverty of cortical sulcation and shallow Sylvian fissure, compatible with cortical developmental malformation like polymicrogyria. Multiple hypointense foci on the susceptibility weighted MRI sequence, diffuse on cortical/subcortical junction, on basal ganglia and, fewer periventricular were seen, some corresponding to the calcifications found on CT. There were other cortical/subcortical areas of T1 hyperintensity, predominantly on temporo-parietal parenchyma, that did not match the ones seen on the CT, which may correspond to hemosiderin or laminar cortical necrosis. Moderate non-hypertensive, asymmetric dilatation of the supratentorial ventricular system, absence of the corpus callosum splenium and asymmetry of the thalamus were also apparent. The posterior fossa was relatively preserved (Figure 4) and asymmetry between the neurocranium and the viscerocranium and retrognathism, were noticeable, 3 months after birth.

**Figure 2.**
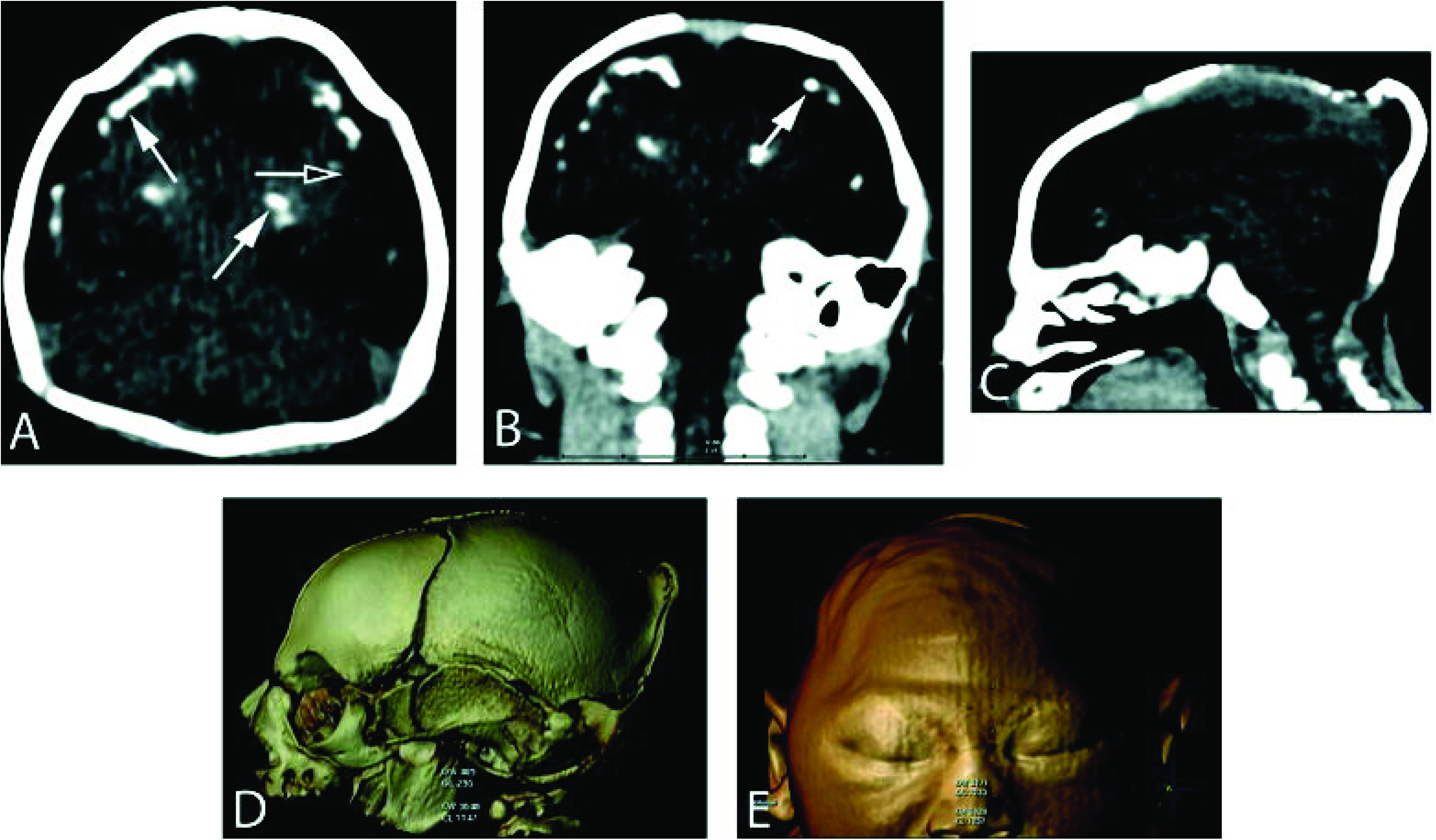
Head CT, obtained at one day of life. Axial (A), coronal (B) and sagittal (C) images show cortical and subcortical, basal ganglia and periventricular calcifications (white arrows); and a simplified gyral pattern and shallow sylvian fissures, compatible with pachygyria (black arrow). Tridimensional (3D) reformatted lateral (D) and frontal (E) images using volume rendering (VR) algorithm demonstrate microcephaly. In D there is an evident skull deformity presenting as occipital protrusion, forming a “spur”, and overriding parieto-occipital bones. In E, notice also the presence of prominent skin folds in high convexity.

**Figure 3.**
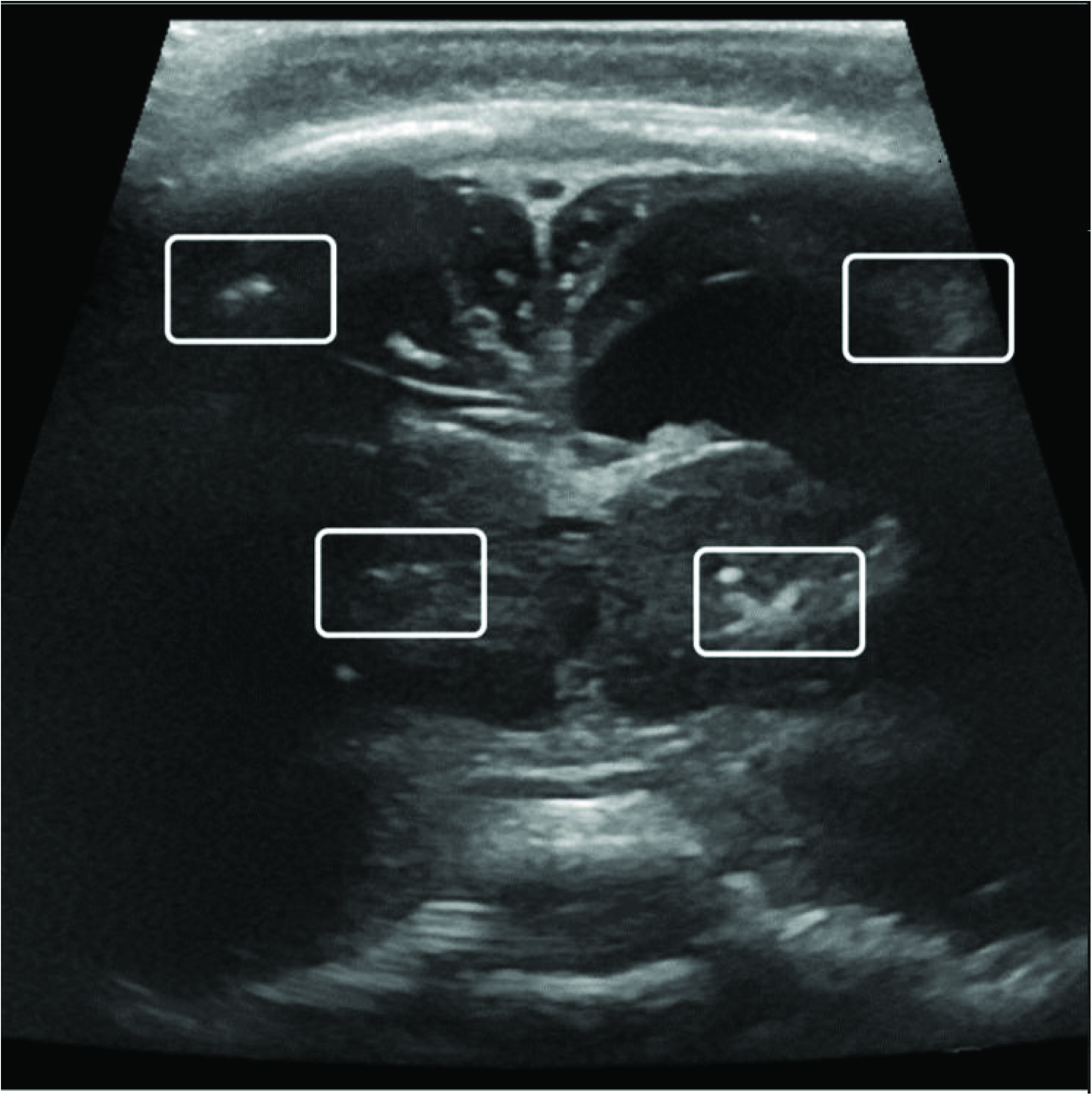
Coronal view of transfontanelar ultrasound. Image obtained at 3 months of life. Notice hyperechogenic foci (rectangles) compatible with the calcifications. There is also ventricular dilation.

**Figure 4.**
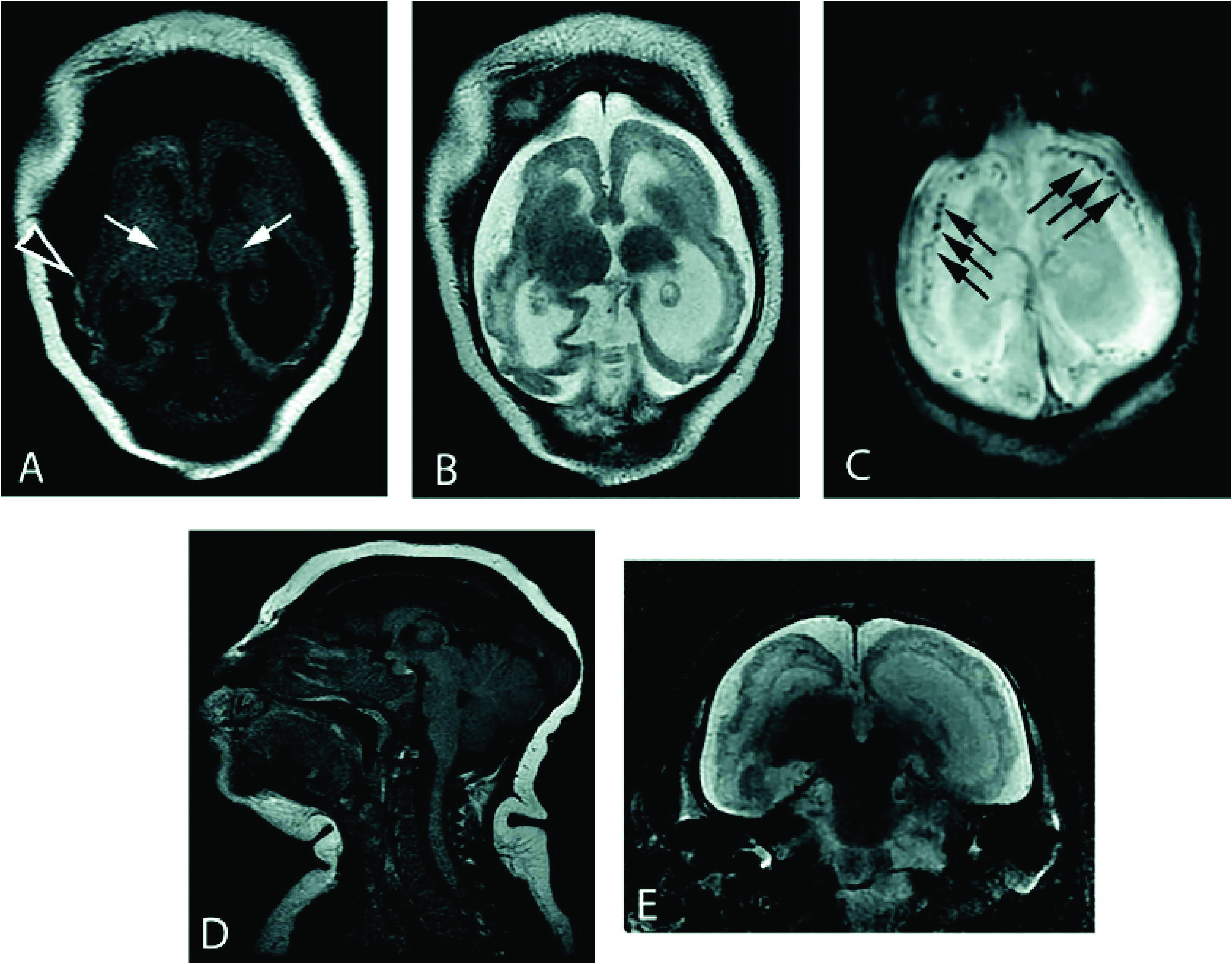
MRI obtained at 3 months of age. Axial T1 (A) and T2-weighted (B) images disclose areas of T1-hyperintensity, which correspond to intracranial calcifications (black arrowhead), in correlation with previous CT. There is clearly a simplified gyral pattern (pachygyria), although in the left temporoparietal region there is an area suggestive of polymicrogyria. Thalamic asymmetry can also be seen, with the left one smaller (white arrows). Axial susceptibility-weighted image (C) shows multiple cortical / subcortical and periventricular foci of low signal intensity, also corresponding to intracranial calcifications (black arrows), again in correlation with CT images. Sagittal T1-weighted image (D) demonstrates craniocephalic disproportion and a relatively preservation of posterior fossa structures. Coronal T2-weighted image (E) discloses asymmetric ventricular dilation, and also the above-mentioned malformations of cortical development, bilaterally. Some hypointense subcortical foci on T2-weighted images may also correspond to calcifications.

### Molecular biology, serology, immunology and virology

Saliva, urine, serum and PBMCs from both mother and child at 3 months of age were negative by PCR for genomic RNA of Chikungunya virus (CHIKV), Yellow fever (YFV), dengue virus (DENV) and Zika virus (ZIKV) (Supplementary Figure 2). Mother’s ELISA results were IgM negative for all flaviviruses and CHIKV. However, the newborn had IgM for ZIKV. Both mother and child were IgG positive for DENV, YFV and ZIKV. We also did plaque neutralization virological tests (PRNT) to confirm the ELISA results. PRNT showed that, the mother sera neutralized 90% of ZIKV up to a 1:80 dilution. Both, the mother and the child had PRNT_90_ for DENV2. Moreover, the mother’s serum had negative PRNT_90_ for West Nile virus (WNV) and YFV. The mother had mean titer values of 137.245 and 48.76 against DENV2-NGC and ZIKV respectively, while the child had only a weak response of 9.635 to DENV2-NGC (Supplementary Table 1 and Supplementary Figure 3). Further serologic confirmation of ZIKV and DENV were done by indirect IF assay on ZIKV infected C6/36 cells with sera from the mother and the child (Figure 5). The clinical records of the mother confirmed previous infections by DENV, CMV, *T. gondii* and syphilis. Therefore, we looked at the immune cytokine profile of the mother PBMCs. The serological response pattern of the mother indicated high amounts of IFN-γ and TNF-α in the supernatants of hers re-stimulated PBMCs (Figure 6) may be a reliable indication of either recent infection, as by ZIKV, or of a higher abundance of memory T cells. Accordingly, this was supported by the increase in IL-2 production, an important cytokine for T cell expansion. Interestingly, these findings are in accordance with data shown in figure 7, showing up to 7.5% of both Th1 and Th17 lymphocytes. This was corroborated by the higher amount of these cytokines in the supernatants when cultures were also stimulated with PMA + Ion, as depicted in figure 6. Moreover, PBMCs also secreted significant amounts of IL-6, but no IL-10, an important anti-inflammatory cytokine. These results are critical because they demonstrate the functional profile of the mother’s T cells, which play a role in controlling viral growth by inducing an antiviral cellular, possibly during Zika infection [16].

**Figure 5.**
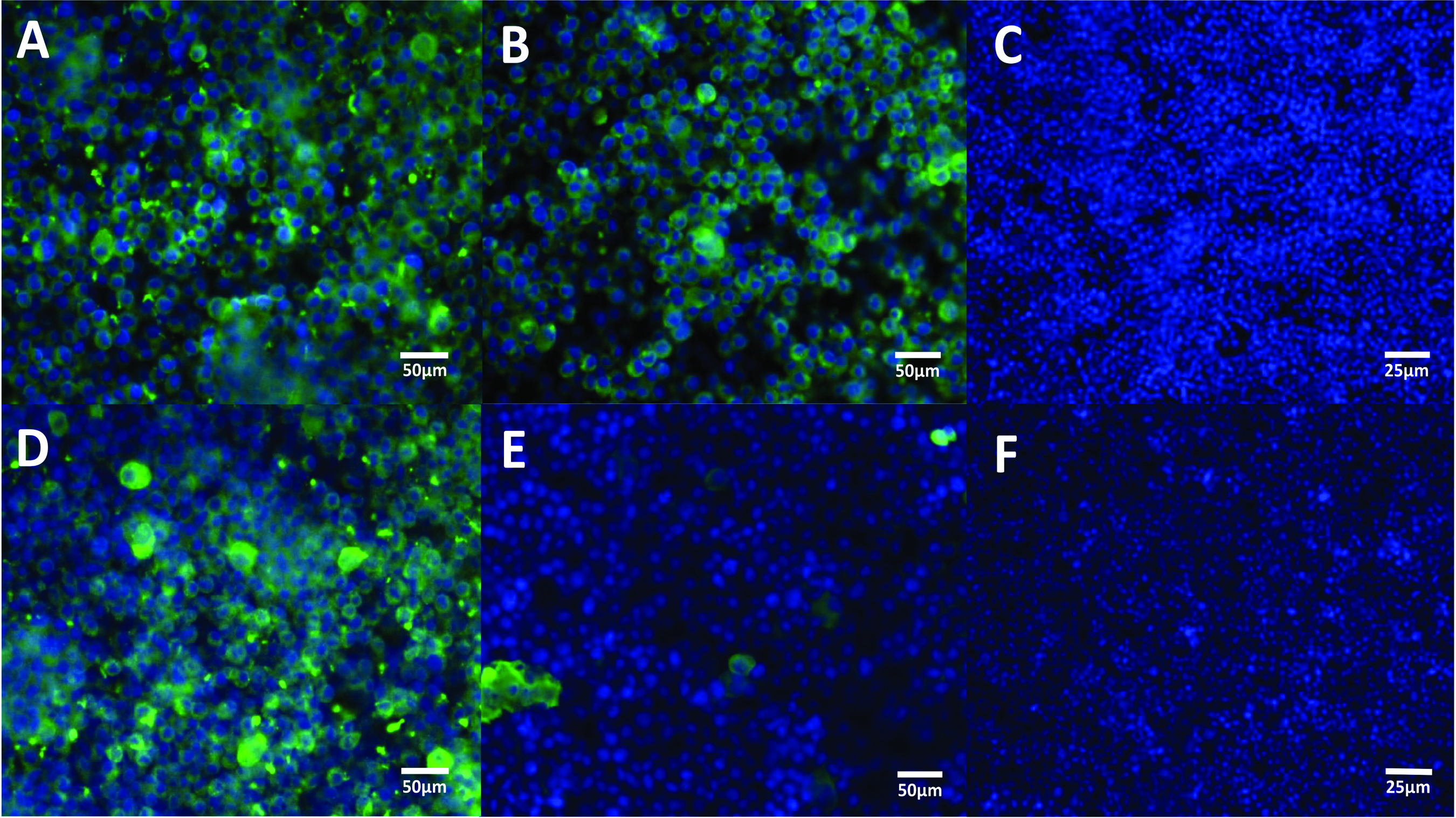
Immunofluorescence of ZIKV and Dengue infection *in vitro*. **A)** C6/36 cells were infected with ZIKV p and exposed to serum of the mother; **B)** exposed to serum of the patient and **C)** exposed to serum from a negative control. **D)** C6/36 cells infected with Dengue virus and exposed to serum of the mother, **E)** exposed to serum of the patient and **F)** exposed to serum from a negative control. Magnifications of 40x (**A**, **B**, **D**, **E**) and 20x (**C**, **F**). Scale bars of 50μm(**A**, **B**, **D**, **E**) and 25 μm (**C**, **F**).

**Figure 6.**
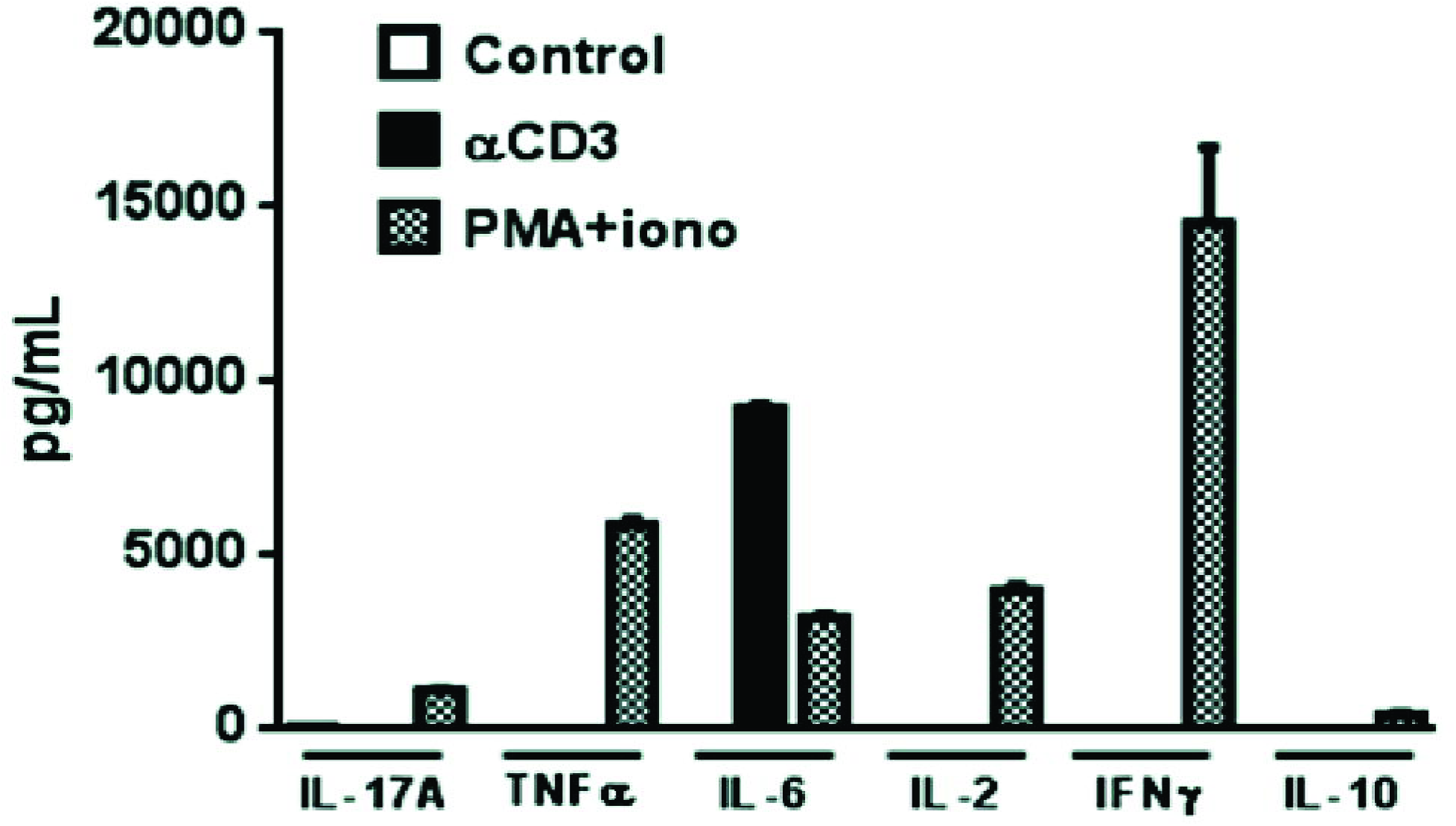
Supernatant cytokine profile of *in vitro* grown the mother’s PBMCs. PBMCs were collected and restimulated in vitro with polyclonal activators anti-CD3 (2 μg/ml) or phorbol-myristate acetate (50 ng/ml) + ionomycin (1 μg/ml). 48 hours later supernatants were collected and submitted to analysis by cytometric bead array for IL-2, IL-6, IL-10, IL-17, IFN-γ and TNF-α according to manufacturer’s protocols. The increased amounts of IFN-γ and TNF-α indicate a Th1 shifted profile, which is consistent with the activation of the pool of memory T cells, including those possibly elicited by the infection DENV or ZIKV.

**Figure 7.**
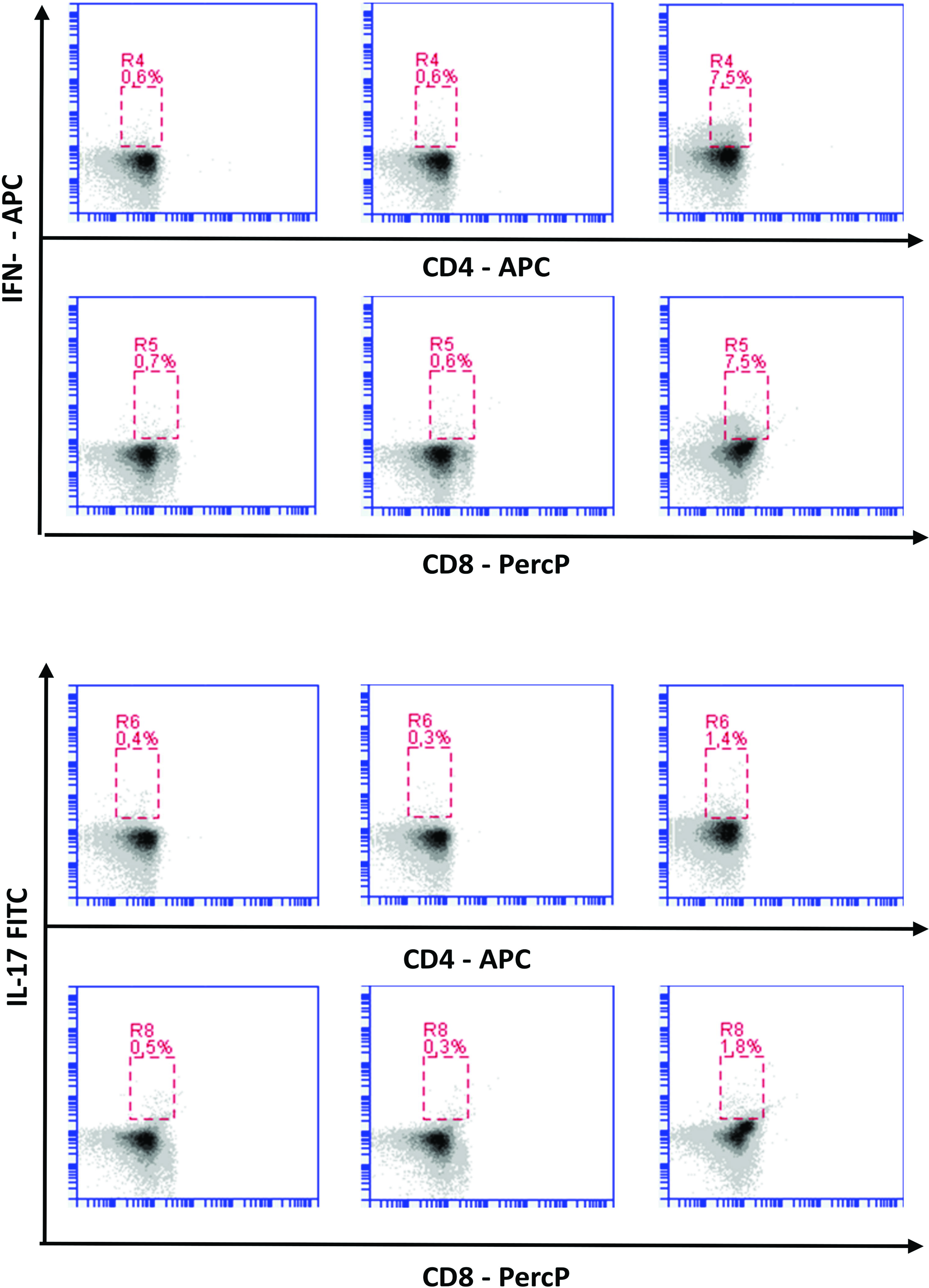
Flow cytometric analysis of the frequency of IFN-g and IL-17-secreting T CD4 and T CD8 lymphocytes from peripheral blood of the mother. PBMCs were obtained and further plated and stimulated or not with anti-CD3 / CD28 (2μg/ml) or PMA (50 ng/ml) + ionomycin (1 μg/ml) overnight in the presence of brefeldin A (1 μg/ml). Cells were further submitted to flow cytometry staining protocols for CD4 APC, CD8 PercP, IFN-□ Pe and IL-17 FITC. The *y*-axis represents cells positive for IFN-γ (top) or IL-17 (bottom) and in the *x*-axis cells positive for CD4 (top) and CD8 (bottom) is shown. Red windows indicate gates used to select double positive cells, *i.e.*, CD4^+^IFN-γ^+^, CD4^+^IL-17^+^, CD8^+^IFN-γ + and CD8^+^IL-17^+^ and numbers indicate the frequency of positive cells. The increase in the frequency of IFN-γ-secreting T CD8 and CD4 lymphocytes corroborates the findings in figure 6 evidencing a shift toward a Th1 immune response.

## Discussion

### Primary microcephaly

Fetal abnormalities were observed in ultrasonography from 12 out of 42 ZIKV-positive mothers (29%) in Rio de Janeiro out of a cohort of 88 mothers with and 88% of seroprevalence for DENV, but no exposure to STORCH agents were reported [19]. Herein we report one of the first cases of microcephaly in the São Paulo - the more densely populated area in Brazil -, shown to be associated with ZIKV by virus seroneutralization test. The definition of microcephaly varies according to age. In the prenatal setting, microcephaly is defined on B-mode ultrasound by an occipitofrontal circumference measurement 2 standard deviations (SD) below the mean for a given age, sex, and gestation [20]. After birth, it is defined as a head circumference three standard deviations (SD) below the reference value for the child’s age [21]. Based on the evidence collected, we would argue that we present a primary microcephaly case, possibly resulting from abnormal brain development related to a developmental insult during the time-specific period (22^nd^ week) of major neural cellular migration, neuronal synaptogenesis and apoptosis [22]. At around the 32^nd^ week brain alterations appear to be noticeable, while the brain stem, cerebellum and posterior fossa, all of which are already well formed do not appear to affected. This was an obvious prenatal intracranial abnormality associated with a clinical picture of an infectious fetopathy as indicated by the serological, immunological and virological data presented.

### Immunological context of pathogen-associated microcephaly

A key finding was the preponderance of IFN-γ, confirming that the mother was indeed exposed to several pathogens that fit our observations of a complex infections profile. Humoral immune responses are well known for their capacity to neutralize viral particles and thus avoid cell invasion. In this context, antibody isotype switching [23] is mainly orchestrated by T helper cells subtypes, in which IFN-γ plays an important role during viral infections. It is noteworthy that cytotoxic T CD8+ lymphocytes are also able to secrete this cytokine, aside from their well known cytotoxic function [24]. Moreover, it is plausible to think that the increase in IFN-γ secreting T cells, may correlate with an increased pool of memory T cells, and less likely to acute responder lymphocytes. The increase in IL-2 secretion observed may also corroborate this possibility. Probably, this seems to be the case as the mother had signs of previous infections by several different pathogens, such as syphilis, *Toxoplasma gondii* and CMV, HSV, DENV and possibly by ZIKV during pregnancy. Altogether, the presence of other pro-inflammatory cytokines, as TNF-α and IL-17, also indicate the status of the adaptive immunological response of the mother that, altogether, may correlate with the dystrophy in the fetal brain tissue, as both Th1 and Th17 cells can invade the central nervous system during neuroinflammation [25]. Besides, raise in IL-6 correlates both, with innate immune activation, as well as with Th17 expansion. On the other hand, IL-10, secreted by suppressive cells and macrophages, is hardly detected [26]. In fact, it has been demonstrated that IFN-γ activates microglial cells in order to restrain toxoplasma spread in the CNS [27] and also to stop CMV replication in mother-to-fetus infection experimental models [28].

### Distinct infectious etiologies of microcephaly

With the exception of rubella, for which she was vaccinated, the mother was at some point exposed or infected by all other STORCH agents (*i.e.*, syphilis, toxoplasmosis, cytomegalovirus and herpes virus), which are pathogens known to cause microcephaly and induce intracerebral calcifications [29]. Congenital syphilis is included in the STORCH agents list, and is known to cause congenital malformations. Herpes virus type 2 infection can undergo transplacental transmission, accounting for only a small fraction of congenital infections, usually associated to chorioretinitis and microcephaly and/or ventricular dilatation [29]. Damage to the placenta can lead to localized inflammation, which could result in localized disruption of the syncytiotrophoblast epithelium, facilitating infection of the fetus. In a cohort study, toxoplasmosis was the only risk factor with significant odds-ratio associated with ZIKV microcephaly in Brazil [30], with a prevalence up to 80% in the South. Although the mother was immunocompetent, had immunity and amniotic fluid PCR negative to *T. gondii*, we observed that the she had a 32-fold increase of IgG titer against toxoplasma during pregnancy. Moreover the child, at 8 days after birth, had even higher titers than those reported for the mother. As a consequence, it is possible that toxoplasma may have had a role in the clinical evolution of this case. It is well known that *T. gondii* associates with the protective epithelium in the placenta and can lead to a local inflammatory response. This process may recruit other immune cells, such as dendritic and placental Hofbauer cells. Fetal placental Hofbauer macrophages (HBCs) are known to be infected by RNA and DNA viruses and may lead to the infection of other placental cells helping the virus reaching the fetus [16,31]. Moreover, the child’s CT scan clinical data was also compatible with severe manifestations caused by congenital CMV and the mother was also seropositive for CMV. DENV has a prevalence that varies from 50% in the Southeast to more than 80% in the Northeast of Brazil. Therefore, the seropositivity for DENV we observed raises a question on the role of heterotypic antibodies produced by previous DENV infection, which could possibly facilitate, or enhance ZIKV infection in the placenta and in the fetus brain. Crucially, the associations we observed to DENV and possibly *T. gondii* were also present in the ZIKV-associated microcephaly case reported by Mlakar et al. [9]. Although a serologic study of 23 infants (out 88) with microcephaly from Pernambuco, did not indicated serologic evidence for TORCH infections [32], 77% of the microcephaly cases observed in Recife are among poor low-income families, already exposed to poor sanitation, poor nutrition and illness, as reported by Brazilian health authorities. On the basis of evidence amassed so far, the Center for Disease Control and Prevention (CDC) has concluded that ZIKV causes microcephaly and other congenital malformations [33]. Nevertheless, given the complex backcloth of STORCH pathogens and DENV infections we report here and given their high prevalence in Brazil, a possible role of STORCH agents in the congenital ZIKV syndrome needs to be addressed at the population level.

## Acknowledgements

This study was approved by an institutional review boards and biological samples were collected under the approval of the Ethics Committee of the Instituto de Ciências Biomédicas (ICB), USP.

---

**Supplementary Figure 1. Serum anti-ZIKV and anti-DENV2 IgG responses measured in the mother and the child.** Columns represent the mean IgG titer values measured with serum samples. Mother serum had anti-DENV2-NGC and anti-ZIKV IgG responses, while child serum had only a minor response against DENV2-NGC.

**Supplementary Figure 2. Viral detection by PCR.** (A) ZIKV Real time PCR results with obtained CTs. (B) Agarose gel showing the amplification pattern of the endogenous control RPL27 from samples after first and second sampling. Negative control consisted of no RNA in the reaction.

**Supplementary figure 3.** Whole body radiography, anteroposterior view. Long bones appear normal, suggestive of no congenital malformation due to syphilis.

## References

1. Dick G, Kitchen S, Haddow A. Zika virus (I). Isolations and serological specificity. Trans R Soc Trop Med Hyg. 1952;46: 509–520.

2. Faye O, Freire CCM, Iamarino A, Faye O, de Oliveira JVC, Diallo M, et al. Molecular Evolution of Zika Virus during Its Emergence in the 20th Century. PLoS Negl Trop Dis. 2014;8: e2636.

3. Lanciotti RS, Kosoy OL, Laven JJ, Velez JO, Lambert AJ, Johnson AJ, et al. Genetic and Serologic Properties of Zika Virus Associated with an Epidemic, Yap State, Micronesia, 2007. Emerg Infect Dis. 2008;14: 1232–1239.

4. Freire CC de M, Iamarino A, Lima Neto DF de, Sall AA, Zanotto PM de A. Spread of the pandemic Zika virus lineage is associated with NS1 codon usage adaptation in humans. bioRxiv. 2015; 1–8.

5. PAHO. Neurological syndrome, congenital malformations, and Zika virus infection. Implications for public health in the Americas. Pan Am Heal Organ. 2015; 1–11.

6. Faria NR, Azevedo R d. S d. S, Kraemer MUG, Souza R, Cunha MS, Hill SC, et al. Zika virus in the Americas: Early epidemiological and genetic findings. Science. 2016;5036: 1– 9.

7. Zanluca C, Campos V, Melo A De, Luiza A, Mosimann P, Igor G, et al. First report of autochthonous transmission of Zika virus in Brazil. Mem Inst Oswaldo Cruz. 2015;110: 569–572.

8. Calvet G, Aguiarv RS, Melo AS, Sampaio SA, de Filippis I, Fabri A, et al. Case Report of detection of Zika virus genome in amniotic fluid of affected fetuses: association with microcephaly outbreak in Brazil. Lancet Infect Dis. 2016;3099: 1–8.

9. Mlakar J, Korva M, Tul N, Popović M, Poljšak-Prijatelj M, Mraz J, et al. Zika Virus Associated with Microcephaly. N Engl J Med. 2016; 1–8.

10. Wagner D, De With K, Huzly D, Hufert F, Weidmann M, Breisinger S, et al. Nosocomial acquisition of dengue. Emerg Infect Dis. 2004;10: 1872–1873.

11. Weidmann M, Faye O, Faye O, Kranaster R, Marx A, Nunes MRT, et al. Improved LNA probe-based assay for the detection of African and South American yellow fever virus strains. J Clin Virol. Elsevier B.V.; 2010;48: 187–192.

12. de Jonge HJM, Fehrmann RSN, de Bont ESJM, Hofstra RMW, Gerbens F, Kamps WA, et al. Evidence based selection of housekeeping genes. PLoS One. 2007;2: 1–5.

13. Sow A, Loucoubar C, Diallo D, Faye O, Ndiaye Y, Senghor CS, et al. Concurrent malaria and arbovirus infections in Kedougou, southeastern Senegal. Malar J. BioMed Central; 2016;15: 1–7.

14. Amorim JH, Porchia BFMM, Balan A, Cavalcante RCM, da Costa SM, de Barcelos Alves AM, et al. Refolded dengue virus type 2 NS1 protein expressed in Escherichia coli preserves structural and immunological properties of the native protein. J Virol Methods. 2010;167: 186–192.

15. Mukherjee S, Dowd KA, Manhart CJ, Ledgerwood JE, Durbin AP, Whitehead SS, et al. Mechanism and significance of cell type-dependent neutralization of flaviviruses. J Virol. 2014;88: 7210–7220.

16. Hamel R, Dejarnac O, Wichit S, Ekchariyawat P, Neyret A, Natthanej L, et al. Biology of Zika Virus Infection in Human Skin Cells. J Virol. 2015;89: 8880–8896.

17. Nelson S, Jost CA, Xu Q, Ess J, Martin JE, Oliphant T, et al. Maturation of West Nile virus modulates sensitivity to antibody-mediated neutralization. PLoS Pathog. 2008;4: e1000060.

18. Salje H, Rodríguez-Barraquer I, Rainwater-Lovett K, Nisalak A, Thaisomboonsuk B, Thomas SJ, et al. Variability in Dengue Titer Estimates from Plaque Reduction Neutralization Tests Poses a Challenge to Epidemiological Studies and Vaccine Development. PLoS Negl Trop Dis. 2014;8: 8–10.

19. Brasil P, Pereira, JP Jr., Raja Gabaglia C, Damasceno L, Wakimoto M, Ribeiro Nogueira RM, et al. Zika Virus Infection in Pregnant Women in Rio de Janeiro — Preliminary Report. N Engl J Med. 2016; 1–11.

20. Salomon LJ, Duyme M, Crequat J, Brodaty G, Talmant C, Fries N, et al. French fetal biometry: Reference equations and comparison with other charts. Ultrasound Obstet Gynecol. 2006;28: 193–198.

21. Krauss MJ, Morrissey AE, Winn HN, Amon E, Leet TL, Hess LW, et al. Microcephaly: An epidemiologic analysis. Am J Obstet Gynecol. 2003;188: 1484–1490.

22. Tau GZ, Peterson BS. Normal development of brain circuits. Neuropsychopharmacology. Nature Publishing Group; 2010;35: 147–168.

23. Geha RS, Rosen FS. The genetic basis of immunoglobulin-class switching. N Engl J Med. 1994;330: 1008 – 1009.

24. Mohr E, Cunningham AF, Toellner K-M, Bobat S, Coughlan RE, Bird R a, et al. IFN-{gamma} produced by CD8 T cells induces T-bet-dependent and-independent class switching in B cells in responses to alum-precipitated protein vaccine. Proc Natl Acad Sci U S A. 2010;107: 17292–17297.

25. Weiner HL. The challenge of multiple sclerosis: How do we cure a chronic heterogeneous disease? Ann Neurol. 2009;65: 239–248.

26. Awasthi A, Carrier Y, Peron JPS, Bettelli E, Kamanaka M, Flavell R a, et al. A dominant function for interleukin 27 in generating interleukin 10-producing anti-inflammatory T cells. Nat Immunol. 2007;8: 1380–1389.

27. Suzuki Y, Sa Q, Gehman M, Ochiai E. Interferon-gamma- and perforin-mediated immune responses for resistance against Toxoplasma gondii in the brain. Expert Rev Mol Med. 2011;13: e31.

28. Bantug GRB, Cekinovic D, Bradford R, Koontz T, Jonjic S, Britt WJ. CD8+ T lymphocytes control murine cytomegalovirus replication in the central nervous system of newborn animals. J Immunol. 2008;181: 2111–2123.

29. Baskin HJ, Hedlund G. Neuroimaging of herpesvirus infections in children. Pediatr Radiol. 2007;37: 949–963.

30. Rocha H, Correia L, Leite A, Campos J, Cavalcante e Silva A, Machado M, et al. Microcephaly: normality parameters and its determinants in northeastern Brazil: a multicentre prospective cohort study. Bull World Heal Organanization. 2016; 1–12.

31. Abbasi M, Kowalewska-Grochowska K, Bahar M a, Kilani RT, Winkler-Lowen B, Guilbert LJ. Infection of placental trophoblasts by Toxoplasma gondii. J Infect Dis. 2003;188: 608– 616.

32. Hazil AN, Poretti A, Cruz DDCS, Tenorio M, Linden A Van Der, Pena LJ, et al. Computed Tomographic Findings in Microcephaly Associated with Zika Virus. N Engl J Med. 2016; 1–3.

33. CDC (2016) CDC Concludes Zika Causes Microcephaly and Other Birth Defects. Available http://www.cdc.gov/media/releases/2016/s0413-zika-microcephaly.html. Accesed 2016 Apr 20.

